# Human Norovirus NS7 Protein Activates Canonical Inflammasome Pathways via NLRP6 in Intestinal Epithelial Cells

**DOI:** 10.1101/2025.11.07.687139

**Authors:** Sunghoon Park, Sung-Gyu Cho, Hyeon Woo Chung, Hwang Su Jin, Taehee Kim, Jae Myun Lee

**Author notes:** These authors contributed equally to this work (Sung-Gyu Cho, Hyeon Woo Chung, Hwang Su Jin, Taehee Kim). Corresponding Author Postal address: 50-1 Yonsei-ro, Seodaemoon-gu, Seoul, 03722, Republic of Korea.

## Abstract

Human norovirus (HNoV) is a leading cause of acute gastroenteritis, yet the mechanisms by which it interfaces with host innate immunity remain elusive. Here, we demonstrate that the HNoV NS7 protein, an RNA-dependent RNA polymerase, acts as a direct activator of canonical inflammasomes. Using reconstituted cell systems and human intestinal enteroids (HIEs), we found that NS7 interacts with both NLRP3 and NLRP6, promoting ASC speck formation, caspase-1 cleavage, and secretion of IL-1β and IL-18. HNoV infection of HIEs recapitulated these events, including gasdermin D processing and robust IL-18 release. Importantly, CRISPR/Cas9-mediated NLRP6 deficiency abrogated inflammasome activation and markedly enhanced viral replication, underscoring the essential role of NLRP6 in epithelial antiviral defense. These findings identify NS7 as a novel inflammasome activator and establish NLRP6 as a key determinant of innate immune control of HNoV. Our study highlights inflammasome signaling as a potential therapeutic target for norovirus infection.

**Author summary:** In this study, we used human intestinal organoid models to explore how norovirus infection triggers a specific immune response known as inflammasome, which helps protect the gut from viral invaders. We focused on a viral protein called NS7 and discovered that it directly activates two types of inflammasome sensors, NLRP3 and NLRP6. We found that NLRP6, which is abundant in the gut lining, is especially important for detecting norovirus and launching an immune response. When we removed NLRP6 from intestinal cells, the virus was able to replicate more easily, and normal immune activation was lost. Our results reveal that norovirus uses its NS7 protein to interact with the body’s immune machinery in the intestine, and that NLRP6 plays a key role in controlling infection. This work highlights a new way in which the gut senses and responds to norovirus and may help guide future efforts to develop treatments that target these immune pathways.

## Introduction

Human norovirus (HNoV) is the leading cause of acute gastroenteritis worldwide, responsible for over 700 million cases and approximately 200,000 deaths annually, particularly affecting children and the elderly(1, 2). Despite its significant public health burden, the mechanisms by which HNoV interacts with host innate immune pathways remain incompletely understood. A key aspect of the innate immune response to viral infection is inflammasome activation, which promotes the maturation and secretion of pro-inflammatory cytokines such as IL-1β and IL-18 and can induce pyroptotic cell death to limit viral spread(3, 4).

Among the various inflammasome-forming pattern recognition receptors (PRRs), NLRP3 is the best characterized and responds to a wide range of viral infections(5). Activation of the NLRP3 inflammasome typically requires a two-step process: a priming signal that induces the transcription of inflammasome components (such as pro-IL-1β and NLRP3 itself), and an activation signal that promotes assembly of the inflammasome complex, leading to caspase-1 activation(6, 7). Several viral proteins have been shown to modulate the NLRP3 inflammasome through direct interactions with its components, either by enhancing or suppressing its activation to favor viral replication or immune evasion(5, 8).

Human norovirus encodes a small number of nonstructural proteins, among which NS7, the viral RNA-dependent RNA polymerase (RdRp), is essential for viral genome replication. Like other viral polymerases, NS7 adopts a 3D^pol^-like structure and contains conserved catalytic motifs that are critical for its enzymatic activity(9, 10). Notably, polymerases from other positive-sense RNA viruses, such as enterovirus 71 and Zika virus, have been reported to modulate NLRP3 inflammasome activation by interacting with NLRP3 and ASC(11, 12). These findings raise the possibility that HNoV NS7 may also function as an immunomodulatory protein during infection.

Furthermore, while most studies of inflammasome activation have focused on immune cells such as macrophages, human norovirus primarily targets intestinal epithelial cells, where the expression profile of inflammasome sensors differs significantly(13, 14). Recent studies have identified NLRP6 as the predominant inflammasome sensor in the intestinal epithelium, where it contributes to antiviral defense and maintenance of mucosal homeostasis(14–16). However, whether HNoV modulates NLRP6 inflammasome activation in epithelial cells has not been investigated.

In this study, we sought to elucidate the role of HNoV NS7 in modulating inflammasome responses. We show that NS7 enhances both NLRP3 and NLRP6 inflammasome activation by interacting with inflammasome components and promoting their assembly. Using a human intestinal enteroid (HIE) model, we further demonstrate that HNoV infection activates the canonical inflammasome pathway in an NLRP6-dependent manner, leading to Caspase-1 activation, Gasdermin D cleavage, and IL-18 secretion. Notably, NLRP6 deficiency results in impaired inflammasome activation and increased viral replication, highlighting its critical role in controlling HNoV infection. These findings provide new insights into how HNoV evades or engages host innate immunity and identify NS7 as a viral factor that modulates inflammasome activity in both immune and epithelial contexts.

## Results

### HNoV NS7 protein enhances NLRP3 inflammasome activation

The human norovirus (HNoV) NS7 protein, an RNA-dependent RNA polymerase (RdRp), is essential for viral genome replication. It adopts a 3D-like polymerase structure with conserved catalytic motifs that are shared among other RNA viruses. Previous studies have shown that similar polymerases from viruses such as Zika and EV71 can modulate NLRP3 inflammasome activation by interacting with inflammasome components, suggesting a mechanism through which viruses may manipulate host immunity.

To examine whether the NS7 protein of HNoV (GII.4 strain) modulates NLRP3 inflammasome activation, we cloned and expressed the NS7 gene in a reconstituted inflammasome system using HEK293T-ASC+CASP1 cells. Co-transfection of Flag-NLRP3 and increasing amounts of NS7-6xMyc plasmid led to dose-dependent cleavage of pro-Caspase-1 into its active p20 subunit, along with increased NS7 expression (Figure 1A). These findings suggest that NS7 enhances NLRP3 inflammasome activation in this system.

**Figure 1.**
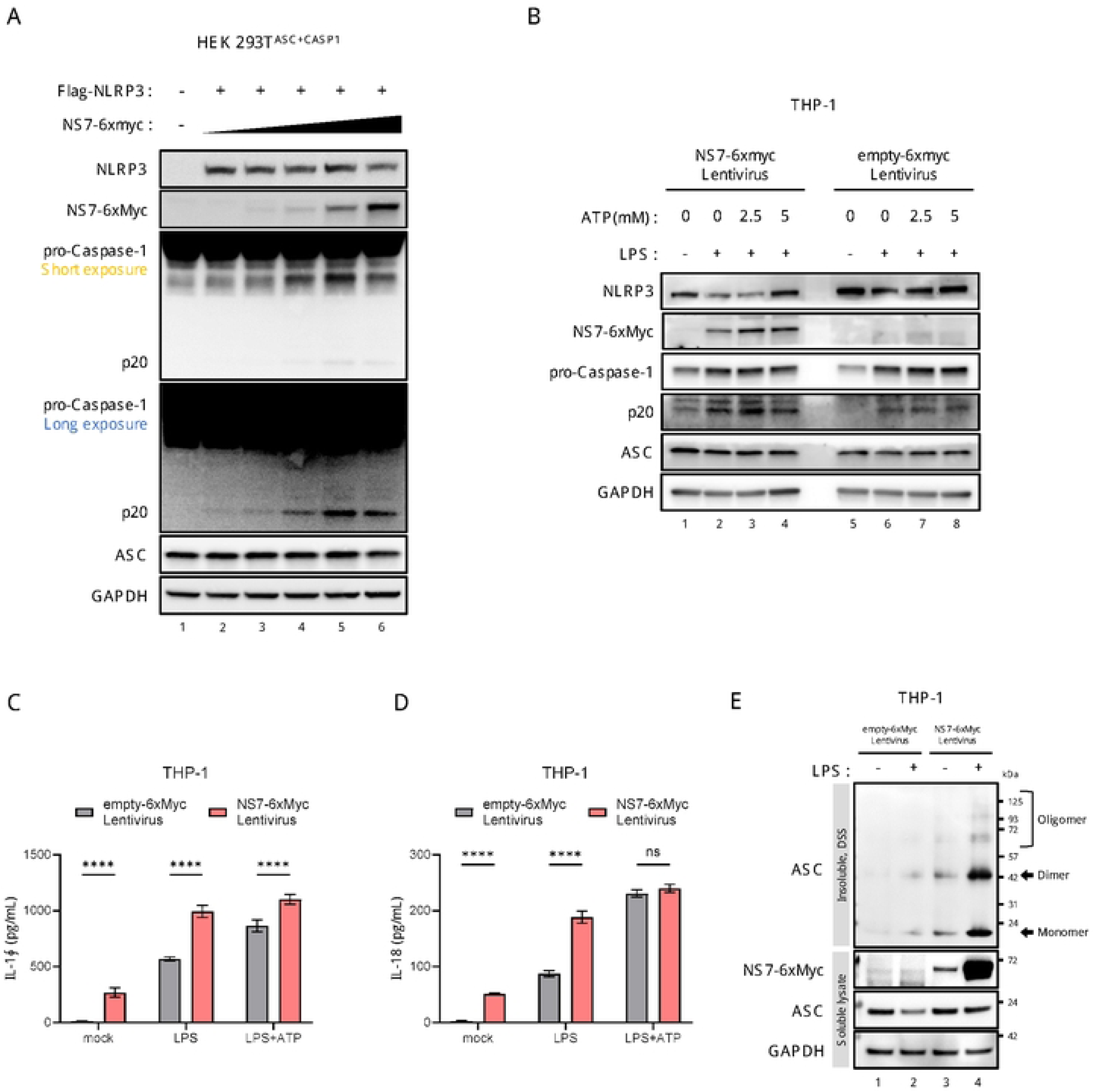
HNoV NS7 protein modulates NLRP3 inflammasome activation in HEK293T and THP-1 cells. (A) HEK293T^ASC+CASP1 cells were transfected with Flag-NLRP3 and increasing amounts of NS7-6xMyc. Cells were harvested 24 hours post-transfection, and protein lysates were analyzed by immunoblotting. Both short and long exposures are shown for pro-Caspase-1 to highlight differences in cleavage efficiency. (B) THP-1 cells transduced with lentivirus expressing NS7-6xMyc or an empty vector (mock) were stimulated with LPS and ATP as indicated. Protein lysates were analyzed by western blotting. (C, D) Supernatants from THP-1 cells treated as in (B) were analyzed for IL-1β (C) and IL-18 (D) secretion using ELISA. (E) THP-1 cells were lysed with 0.5% NP-40 lysis buffer, and insoluble pellets were crosslinked with DSS. Western blotting was performed to detect ASC oligomerization. Soluble fractions were also analyzed for NS7-6xMyc, ASC, and GAPDH expression.

We further investigated the role of NS7 in THP-1 cells. Lentiviral transduction was used to stably express NS7-6xMyc or an empty vector (mock control). Following LPS priming and ATP stimulation, NS7 expression led to enhanced Caspase-1 cleavage compared to controls (Figure 1B). ELISA analysis revealed significantly elevated secretion of IL-1β and IL-18 in NS7-expressing cells (Figure 1C, D). ASC oligomerization, a hallmark of inflammasome assembly, was also confirmed via DSS-crosslinking of insoluble pellets (Figure 1E). Together, these results indicate that HNoV NS7 promotes NLRP3 inflammasome activation by enhancing Caspase-1 cleavage, cytokine secretion, and ASC oligomerization, supporting its role as a potential modulator of innate immune responses.

### NS7 protein interacts with NLRP3 inflammasome components

Given that NS7 enhances NLRP3 inflammasome activation, we hypothesized that it might act as a pathogen-associated molecular pattern (PAMP) facilitating inflammasome assembly. Co-immunoprecipitation (co-IP) in HEK293T cells co-transfected with Flag-NLRP3 and NS7-6xMyc plasmids confirmed a direct interaction between NS7 and NLRP3 (Figure 2A).

**Figure 2.**
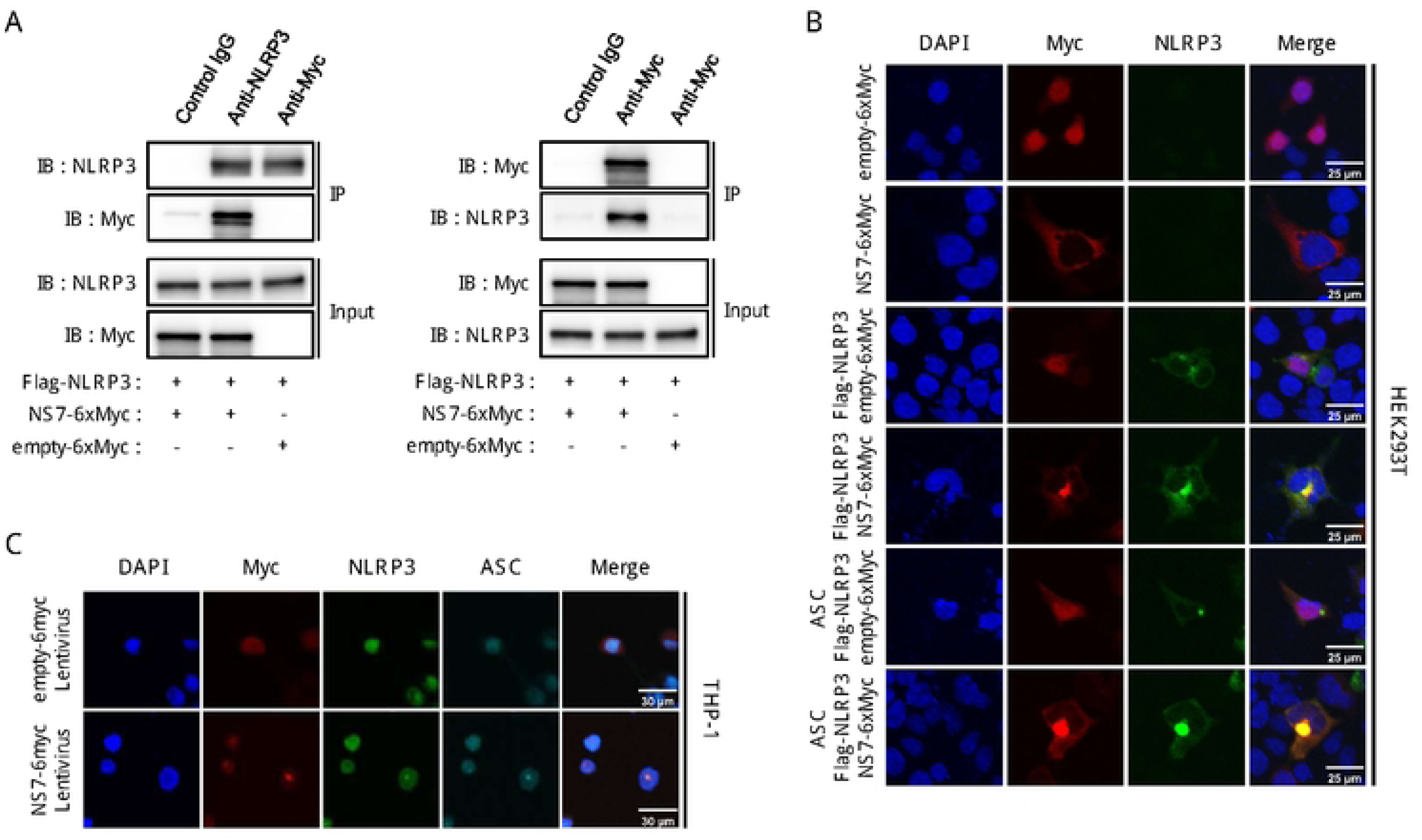
NS7 protein interacts with NLRP3 and co-localizes with inflammasome components. (A) HEK293T cells were transfected with Flag-NLRP3, NS7-6xMyc, or empty-6xMyc plasmids as indicated. Co-immunoprecipitation (co-IP) was performed using anti-NLRP3 or anti-Myc antibodies, followed by immunoblotting (IB). Input lysates were also analyzed for NLRP3 and NS7-6xMyc expression. (B) Confocal microscopy images of HEK293T cells co-transfected with Flag-NLRP3, NS7-6xMyc, and ASC plasmids. Co-localization of NLRP3 (green), NS7-6xMyc (red), and nuclei (DAPI, blue) is shown. Scale bar: 25 μm. (C) Confocal microscopy analysis of THP-1 cells transduced with lentivirus expressing NS7-6xMyc or an empty vector and stimulated with LPS. Co-localization of NLRP3 (green), ASC (cyan), NS7-6xMyc (red), and nuclei (DAPI, blue) is shown. Scale bar: 30 μm.

Confocal microscopy of HEK293T cells co-expressing ASC revealed cytoplasmic co-localization of NS7 and NLRP3 (Figure 2B). In PMA-differentiated THP-1 cells, NS7 expression alone was sufficient to induce inflammasome assembly without additional stimulation, as demonstrated by the co-localization of NS7-6xMyc, NLRP3, and ASC (Figure 2C). These data indicate that NS7 directly interacts with key inflammasome components and promotes their assembly in both HEK293T and THP-1 cells.

### NS7 protein modulates NLRP6 inflammasome activation

Since HNoV primarily infects intestinal epithelial cells, where NLRP3 expression is minimal, we next focused on NLRP6, which is the dominant inflammasome sensor in these tissues. Re-analysis of public RNA-seq datasets from human duodenum, jejunum, and ileum confirmed that NLRP6 is highly expressed, whereas NLRP3 and other NOD-like receptors were largely absent (Figure 3A). Western blot analysis of human intestinal enteroid (HIE) lysates corroborated these findings, showing robust expression of NLRP6 but undetectable levels of AIM2, NLRP3, and NLRC4 proteins (Figure 3B).

**Figure 3.**
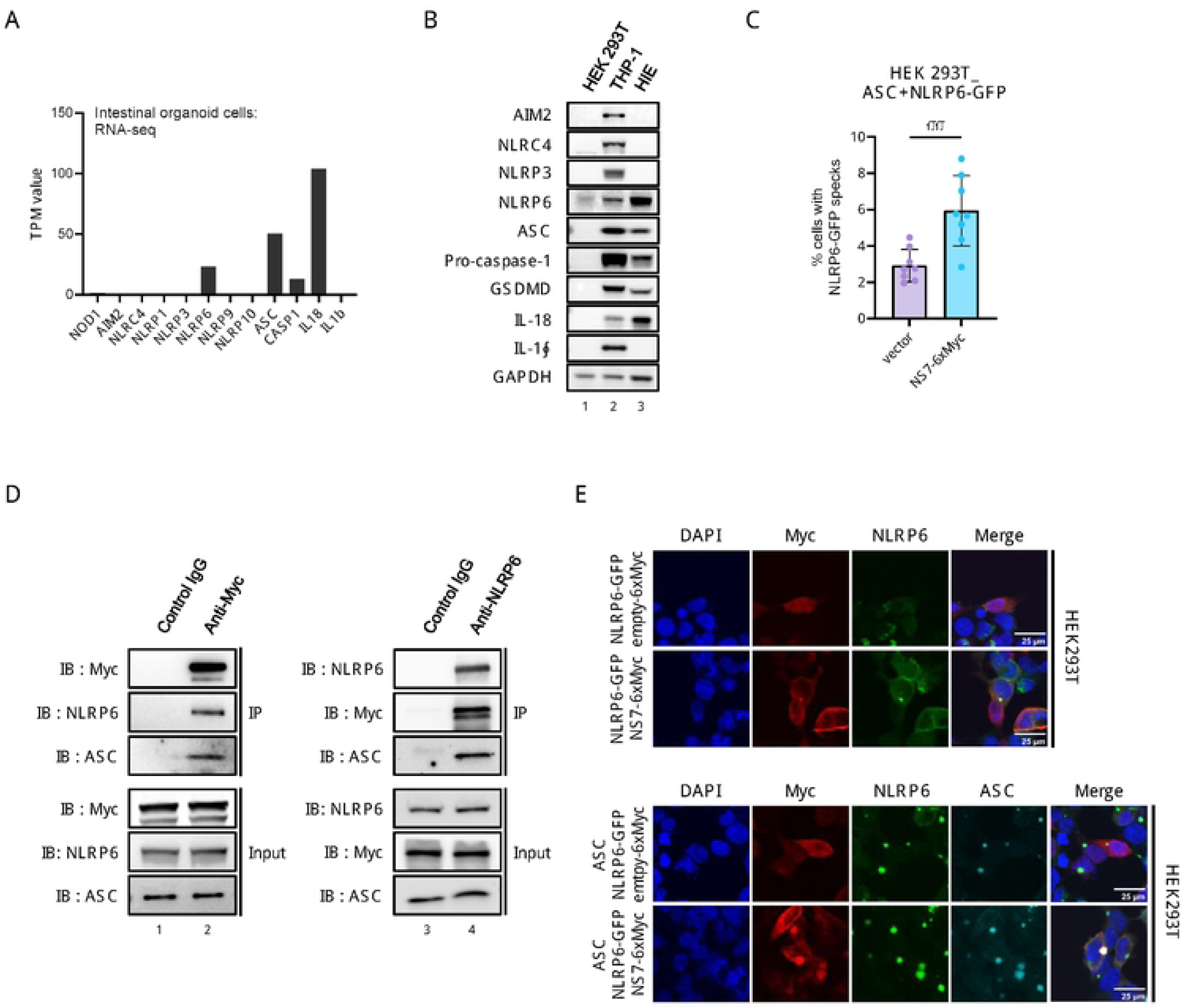
NS7 protein interacts with NLRP6 inflammasome components and promotes NLRP6 speck assembly. (A) RNA-seq analysis of human intestinal enteroids (HIEs) from the duodenum, jejunum, and ileum. TPM values of inflammasome components were compared to assess their expression profiles. (B) Western blot analysis of inflammasome-related proteins in HEK293T cells, THP-1 cells, and HIE lysates to confirm protein-level expression. (C) Quantification of NLRP6-GFP specks in HEK293T cells co-transfected with ASC, NLRP6-GFP, and either NS7-6xMyc or empty-6xMyc plasmids. Data are presented as mean ± SEM from three independent experiments. (D) Co-immunoprecipitation (co-IP) analysis of HEK293T cells co-transfected with NLRP6-GFP, NS7-6xMyc, and ASC, assessing the interaction between NS7 and NLRP6. (E) Confocal microscopy images of HEK293T cells co-transfected with NLRP6-GFP, NS7-6xMyc, and ASC. Co-localization of NS7-6xMyc (red), NLRP6-GFP (green), ASC (cyan), and nuclei (DAPI, blue) is shown. Scale bar: 25 μm.

Notably, NS7 co-expression led to a significant increase in the number of cells exhibiting NLRP6 specks (Figure 3C). Given the observation, we next examined whether NS7 interacts directly with NLRP6. To address this, we performed co-immunoprecipitation experiments in HEK293T cells co-transfected with NLRP6-GFP, ASC, and NS7-6xMyc. These assays confirmed a direct interaction between NS7 and the NLRP6 inflammasome complex (Figure 3D). Furthermore, confocal imaging revealed cytoplasmic co-localization of NS7 and NLRP6 (Figure 3E). Collectively, these results demonstrate that NS7 enhances NLRP6 inflammasome activation, likely through direct interaction and spatial coordination with key inflammasome components.

### Human norovirus infection induces inflammasome activation in human intestinal enteroids

To determine whether HNoV infection triggers inflammasome activation in HIEs, we infected HIEs with the GII.4 strain and monitored viral replication and inflammasome activity. qRT-PCR analysis confirmed robust viral replication, with increasing genome equivalents detected at 1, 48, and 96 hours post-infection (hpi) compared to mock-infected controls (Figure 4A).

**Figure 4.**
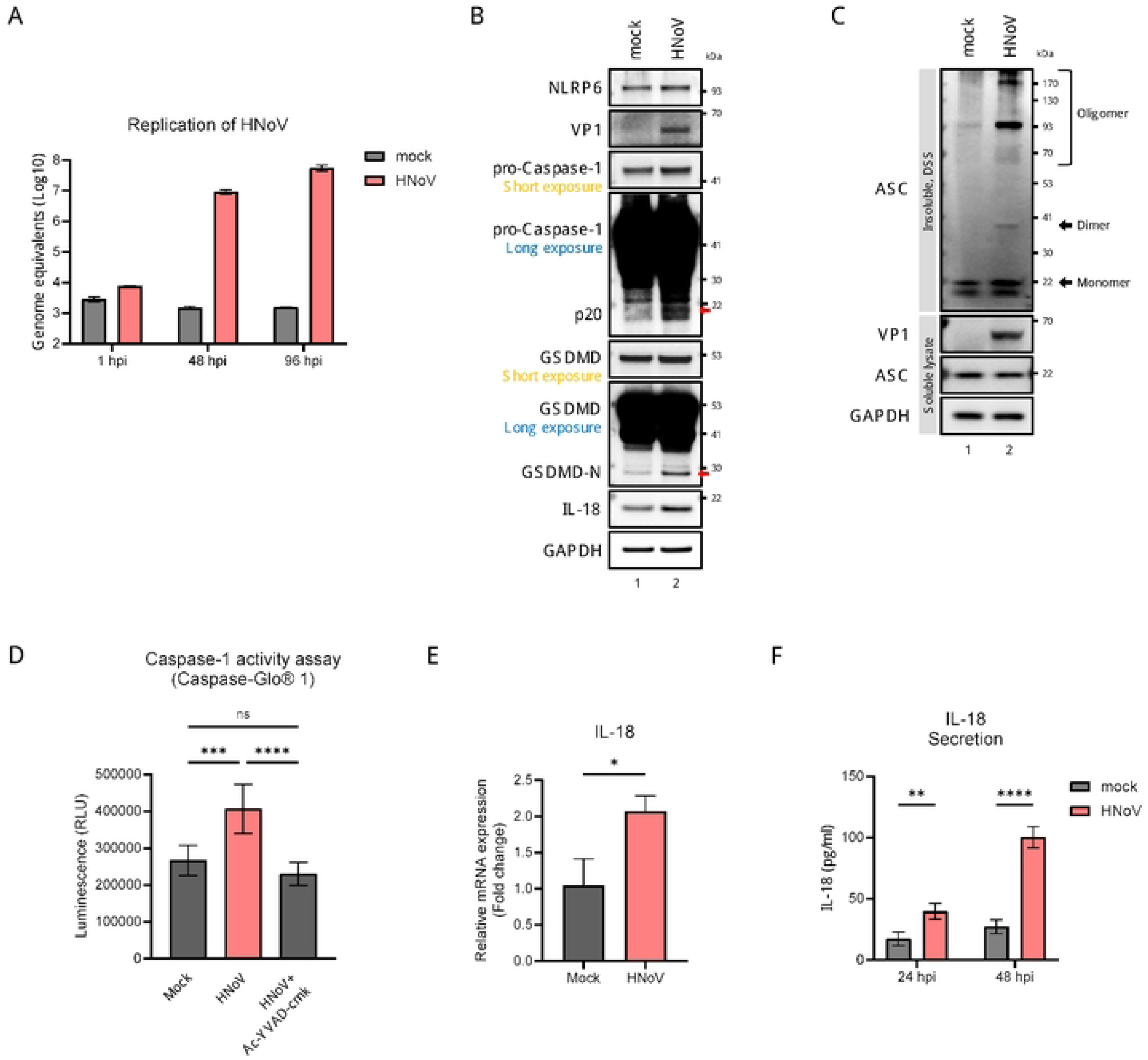
HNoV infection induces inflammasome activation in human intestinal enteroids (HIEs). (A) Quantification of HNoV replication in HIEs by qRT-PCR. HIEs were infected with GII.4 stool filtrate (1 × 10⁷ GE/mL), and viral RNA levels were measured at 1, 48, and 96 hpi. Mock infection was performed using heat-inactivated virus. (B) Caspase-1 activity in HIEs at 48 hpi, measured using the Caspase-Glo® 1 assay. Luminescence was recorded after a 1-hour substrate incubation. Ac-YVAD-cmk (5 μM) was added as a caspase-1 inhibitor. (C) Western blot analysis of lysates from HIEs harvested at 48 hpi. (D) HIEs were lysed at 48 hpi using 0.5% NP-40 buffer. Insoluble pellets were crosslinked with DSS, and western blotting was used to detect ASC oligomerization. Soluble fractions were also analyzed for ASC and GAPDH expression. (E) IL-18 mRNA expression at 48 hpi measured by qRT-PCR. Data were normalized to GAPDH and expressed as fold change relative to mock-infected cells. (F) IL-18 secretion in supernatants at 24 and 48 hpi measured by ELISA. Data are presented as mean ± SEM from three independent experiments (**p < 0.01; ****p < 0.0001; ns, not significant).

Caspase-1 activity, measured at 48 hpi using the Caspase-Glo® 1 assay, was significantly increased in infected cells, and was suppressed by the Caspase-1 inhibitor Ac-YVAD-cmk (Figure 4B). Western blotting showed cleavage of pro-Caspase-1 into its active p20 form and Gasdermin D into its N-terminal fragment (GSDMD-N), which mediates pyroptosis (Figure 4C). IL-18 levels were also increased in infected cells.

ASC oligomerization, indicative of inflammasome assembly, was confirmed in HNoV-infected cells using DSS-crosslinking (Figure 4D). This, together with Caspase-1 activation and GSDMD cleavage, indicates the activation of the canonical ASC-dependent inflammasome pathway. Additionally, qRT-PCR showed increased IL-18 mRNA expression at 48 hpi (Figure 4E), and ELISA confirmed elevated IL-18 secretion at 24 and 48 hpi (Figure 4F). These findings collectively demonstrate that HNoV infection activates canonical inflammasome signaling in human intestinal epithelial cells.

### NLRP6 is required for HNoV-induced inflammasome activation

To elucidate the role of NLRP6 in HNoV-induced inflammasome activation, we generated NLRP6 knockout (KO) HIEs using lentiviral CRISPR-Cas9 transduction. Upon HNoV infection, control HIEs (Lenti-CTL) displayed clear evidence of inflammasome activation—including Caspase-1 and GSDMD cleavage and IL-18 upregulation—whereas these responses were abolished in NLRP6 KO HIEs (Figure 5A). Caspase-1 activity assays further confirmed that NLRP6 was required for inflammasome activation in this context (Figure 5B).

**Figure 5.**
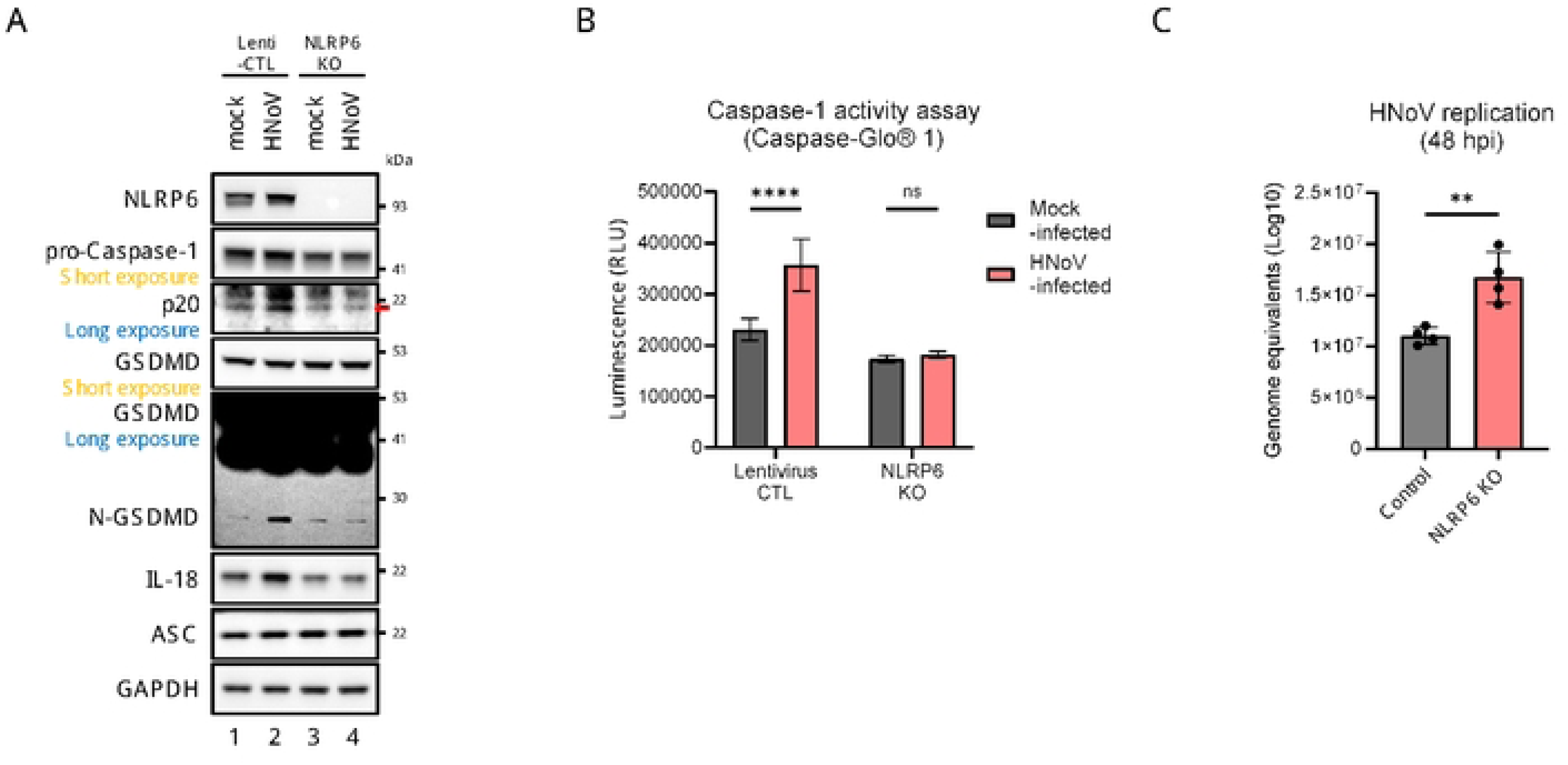
NLRP6 is required for HNoV-induced inflammasome activation in HIEs. (A) Western blot analysis of HIEs transduced with lentivirus expressing Cas9 control (Lenti-CTL) or NLRP6-targeting gRNA (NLRP6 KO). Cells were infected with GII.4 (1 × 10⁷ GE/mL) or heat-inactivated GII.4 and harvested at 48 hpi. (B) Caspase-1 activity in HIEs transduced with Lenti-CTL or NLRP6 KO and infected as in (A). Luminescence was measured at 48 hpi using the Caspase-Glo® 1 assay. (C) Quantification of HNoV replication in NLRP6 KO and Lenti-CTL HIEs by qRT-PCR at 48 hpi. Data are presented as mean ± SEM from three independent experiments (****p < 0.0001; ns, not significant; unpaired t-test).

Strikingly, viral replication was significantly elevated in NLRP6 KO cells at 48 hpi, as assessed by viral genome quantification (Figure 5C). These results indicate that NLRP6 not only mediates inflammasome activation in HIEs but also plays a protective role in limiting HNoV replication. Whether this antiviral effect is mediated solely by inflammasome activation or involves additional pathways remains to be elucidated.

## Discussion

The development of human intestinal enteroid (HIE) models has revolutionized human norovirus (HNoV) research, overcoming a decades-long barrier to studying viral replication and pathogenesis(17, 18). This cultivation system has enabled critical advancements in antiviral and vaccine development, replacing surrogate assays—such as histo-blood group antigen (HBGA) blocking tests—with direct neutralization testing in physiologically relevant models(19). HIEs allow precise evaluation of strain-specific antibody responses and viral inhibition, bypassing the need to infer protection from indirect markers. Beyond their utility in therapeutics, HIEs have provided unprecedented insight into host-pathogen interactions, particularly in innate immune responses. Previous studies using HIEs have shown robust activation of type III interferons (IFN-λ) during HNoV infection. Genetic ablation of STAT1 or pharmacologic inhibition of the JAK/STAT pathway enhances HNoV replication, confirming the importance of interferon signaling in antiviral defense(20–22).

Notably, recent studies have shown that the HNoV NS7 protein undergoes liquid-liquid phase separation (LLPS) during infection, forming biomolecular condensates that function as replication centers(23). These compartments concentrate viral components and create microenvironments optimized for genome replication(24). This property of NS7 adds a new dimension to our understanding of how viral polymerases may interface with host immune systems during infection.

In this study, we demonstrate that the human norovirus NS7 protein, a 3D^pol^-like RNA-dependent RNA polymerase, directly engages both NLRP3 and NLRP6 inflammasomes, thereby linking viral replication machinery to host innate immunity. In vitro, NS7 promoted NLRP3 inflammasome assembly, caspase-1 activation, and IL-1β/IL-18 secretion. These effects mirror those observed in other RNA viruses such as Zika virus and Senecavirus A, whose polymerases similarly modulate inflammasome activation. In human intestinal enteroids, HNoV infection led to robust activation of the inflammasome pathway, evidenced by caspase-1 cleavage, gasdermin D processing, and IL-18 secretion. The detection of ASC oligomerization further confirmed the activation of the canonical inflammasome pathway, as ASC specks form the structural scaffold necessary for caspase-1 recruitment and downstream inflammatory signaling.

Crucially, these inflammasome responses were abolished in NLRP6-deficient HIEs, whereas viral replication was significantly enhanced. These observations identify NLRP6 as a central regulator of canonical inflammasome activation during HNoV infection and highlight its dual role in limiting viral replication and mediating inflammation. This finding aligns with previous studies demonstrating the importance of NLRP6 in intestinal immunity, particularly in coordinating IL-18-mediated antiviral signaling and maintaining epithelial integrity. However, our data also revealed a potential paradox: while NLRP6 activation restricts HNoV replication, it may simultaneously contribute to intestinal inflammation.

Our findings substantially expand the current understanding of NLRP6’s role in the intestinal epithelium during viral infection. The dual role of NLRP6—as both an antiviral effector and a driver of inflammation— underscores the delicate balance that the host must maintain to control infection without inducing excessive tissue damage. The enhanced viral replication observed in NLRP6-deficient HIEs suggests that NLRP6 acts as a frontline defense against HNoV, potentially through multiple mechanisms: pyroptosis of infected cells, IL-18– mediated enhancement of epithelial barrier function, and possibly inflammasome-independent suppression of viral replication complexes. Our findings also bridge the gap between NLRP6’s previously established role in bacterial pathogen defense and its relatively unexplored function in human enteric viral infections.

By demonstrating NLRP6-dependent inflammasome activation in response to HNoV infection, this study provides a molecular framework for understanding how the intestinal epithelium detects and responds to viral pathogens. The identification of canonical inflammasome features—such as ASC oligomerization and caspase-1 activation—in intestinal epithelial cells broadens the known spectrum of antiviral mechanisms in the gut. These insights carry important therapeutic implications, particularly in guiding whether to suppress or preserve inflammatory responses during HNoV infection.

Nevertheless, this study has several limitations. Although HIEs provide a physiologically relevant model, they do not fully replicate the complexity of the intestinal microenvironment, which includes mucus layers, microbiota, and immune cell interactions. In murine models, NLRP6 is known to regulate microbiome composition and to enhance IFN-λ production—both of which contribute to enteric viral suppression(25–27). The absence of these factors in our system complicates the interpretation of enhanced viral replication observed in NLRP6-deficient HIEs. Thus, it remains unclear whether this phenotype results solely from impaired inflammasome activation or also involves inflammasome-independent functions of NLRP6. This ambiguity highlights the multifunctional nature of NLRP6 and the need for further studies to elucidate how its different signaling roles intersect in the gut microenvironment.

In conclusion, our study demonstrated that HNoV infection activates the canonical inflammasome pathway in human intestinal epithelial cells, with NLRP6 acting as a critical regulator of this response. We show that the NS7 protein promotes inflammasome activation by directly engaging inflammasome components, providing a mechanistic link between viral replication and host innate immunity. The disruption of inflammasome signaling, particularly through NLRP6 deficiency, not only enhances viral replication but also impairs immune activation, emphasizing the importance of inflammasome-mediated antiviral defense in the gut. These findings reveal a complex interplay between antiviral immunity and inflammation and suggest that therapeutic strategies targeting the inflammasome may offer new avenues to balance viral control and immunopathology during norovirus infection.

## Materials and Methods

### Ethics statements

Human norovirus samples and intestinal tissue were obtained from patients at Severance Hospital, Seoul with institutional approval. The stool specimen collection was approved by the Institutional Review Board of Severance Hospital, Seoul (IRB No: 4-2020-0480), and written informed consent was obtained from the patient prior to collection. The generation and use of human intestinal enteroids derived from intestinal tissue biopsies were approved by the Institutional Review Board of Severance Hospital, Seoul (IRB No: 4-2012-0859). Written informed consent was obtained from the tissue donor. All procedures involving human-derived specimens and tissues were conducted in accordance with the approvals granted by the Institutional Review Board.

### Cell culture

HEK293T cells were cultured in DMEM with 10% FBS and 1% penicillin/streptomycin at 37°C, passaged every 3–4 days at 80–90% confluency. THP-1 cells were maintained in RPMI 1640 with 10% FBS and 1% penicillin/streptomycin, subcultured to keep 2–5×10⁵ cells/mL, and differentiated with 50 nM PMA for 24 h. Human intestinal enteroids (HIEs) were established from healthy adult ileal crypts isolated by endoscopy, embedded in Matrigel (Corning), and cultured in CMGF+ medium, which consists of advanced DMEM/F12 (Gibco) mixed 1:1 with L-WRN medium and supplemented with B27, N2, N-acetylcysteine, hEGF, [Leu15]-gastrin I, A83-01, SB202190, and Y-27632. Medium was refreshed every 2–3 days. For differentiation, enteroids were dissociated into single cells with TrypLE Express (Gibco), seeded in CMGF+ with Y-27632 for 24 hours to establish monolayers, and then switched to differentiation medium lacking R-spondin and Wnt but containing reduced Noggin (5 ng/mL) for 3–5 days, with medium changes every 1–2 days.

### Isolation of HNoV GII.4

A human norovirus GII.4 Sydney stool specimen was collected from a patient diagnosed with GII.4 infection at Severance Hospital, Seoul, following approval from the Institutional Review Board (IRB No. 4-2020-0480). The stool sample was suspended in phosphate-buffered saline (PBS) at a 1:10 ratio and thoroughly mixed. To remove particulate matter, the suspension was clarified by centrifugation and then sequentially filtered through membranes with pore sizes of 1.0, 0.8, 0.45, and 0.22 μm. The viral titer in the filtered sample was determined by RT-qPCR. A standard curve was generated using a positive control RNA transcript corresponding to the human norovirus capsid region, which was prepared according to the method described by Lee et al. (2011)(28).

### Cloning of HNoV NS7 Gene

Viral RNA was extracted from stool samples positive for human norovirus GII.4/P31 strain. The NS7 (RdRp) gene was amplified via RT-PCR and sequenced to confirm fidelity. For cloning, the PCR product was first inserted into the pDONR™ vector using the BP Clonase™ II enzyme mix (Invitrogen) through a BP recombination reaction. The resulting entry clone (pENTR-NS7) was then recombined into the destination vector pDEST1027-6xMyc using LR Clonase™ II enzyme mix (invitrogen), generating the final expression construct pDEST1027-NS7-6xMyc.

### Immunoblotting

Samples were lysed in 1% NP-40 lysis buffer with protease inhibitor cocktail (GenDEPOT). After lysis, the soluble fraction was collected and protein concentration was determined using the BCA assay (Thermo Fisher Scientific). The insoluble pellet was subjected to crosslinking with DSS for 30 minutes to detect ASC oligomerization, then resuspended in SDS sample buffer. Soluble protein and the crosslinked insoluble fraction were separated on 4–20% gradient SDS-PAGE gels and transferred to nitrocellulose membranes (Invitrogen). Membranes were blocked with 5% skim milk in PBS-T and incubated with primary antibodies against caspase-1 (ab207802; Abcam), ASC (sc-271054; Santa Cruz Biotechnology), NLRP3 (#13158; Cell Signaling Technology), Myc-Tag (#2276; Cell Signaling Technology), GSDMD (#36425; Cell Signaling Technology), IL-18(#54943; Cell Signaling Technology), GAPDH (60004-1-Ig; Proteintech), and NLRP6 (GTX85157; GeneTex). After incubation with HRP-conjugated secondary antibodies, membranes were visualized using a luminescent image analyzer (LAS-4000; FUJIFILM).

### Confocal microscopy

Cells grown on chamber slides were fixed with 4% paraformaldehyde for 15 minutes at room temperature. After fixation, cells were permeabilized and blocked in PBS containing 1% bovine serum albumin (BSA), 5% normal goat serum, and 0.15% Triton X-100 for 1 hour at room temperature. Primary antibodies were applied for 1 hour at room temperature. Cells were then incubated with fluorescent-conjugated secondary antibodies for 1 hour at room temperature in the dark. Nuclei were stained and mounted using VECTASHIELD with DAPI(H-1200-10; Vector Laboratories). Imaging was performed using a confocal laser scanning microscope (FV1000; Olympus) under identical settings. Image analysis were performed using ImageJ software.

### GII.4 HNoV infection of HIE monolayer

For infection of human intestinal enteroid (HIE) monolayers with GII.4 human norovirus, the GII.4-positive stool filtrate was diluted in basal medium (CMGF-) to a final concentration of 1.5×107 GEs/mL. HIE monolayers that had been differentiated for three days were washed twice with CMGF-. The diluted stool inoculum was then added to the washed monolayers and incubated at 37℃ for 5 minutes. Following this, the plates were centrifuged at over 200 × g for 1 hour at temperatures above 32°C. After centrifugation, the cells were washed twice with CMGF-to remove unbound virus and then overlaid with differentiation medium. For analysis of inflammasome activation, cells were harvested 48 hours post-infection.

### Co-immunoprecipitation assays

Whole-cell lysates were prepared by lysis in NP-40 buffer supplemented with protease inhibitors. After clarification by centrifugation, protein concentration was determined, and equal amounts of lysate were used for each immunoprecipitation reaction. Lysates were incubated with control mouse or rabbit immunoglobulin G (IgG; Cell Signaling Technology) or with anti-Myc, anti-NLRP3, or anti-NLRP6 antibodies, together with Dynabeads Protein G (Invitrogen), under gentle rotation at 4°C for 2–4 hours. Beads were washed with cold lysis buffer. Immunocomplexes were eluted by boiling in SDS sample buffer and analyzed by SDS-PAGE followed by immunoblotting.

### RNA extraction and RT-qPCR

Viral RNA was extracted from clarified stool supernatants using the QIAamp Viral RNA Mini Kit (Qiagen) following the manufacturer’s protocol. Norovirus GII RNA levels were quantified by RT-qPCR with the following primers and probe: QNIF2d (5’-ATGTTCAGRTGGATGAGRTTCTCWGA-3’), COG2R (5’-TCGACGCCATCTTCATTCACA-3’), and QNIFS (5’-FAM-AGCACGTGGGAGGGCGATCG-TAMRA-3’)(29). For RNA extraction from HIE cells, the Arcturus PicoPure RNA Isolation Kit (Applied Biosystems) was used. RT-qPCR for norovirus detection used the same primer/probe set, while TaqMan assays (Thermo Fisher Scientific) quantified NLRP6 (Mm00460229_m1), IL-18 (Hs01038788_m1), and GAPDH (Hs02758991_g1) transcripts as an internal control. Thermocycling was performed at 55°C for 10 min, 95°C for 1 min, then 40 cycles of 95°C for 10 s and 60°C for 30 s. Relative expression was calculated by the ΔΔCt method using GAPDH as control.

### Comparative transcriptomic analysis of HIE

Transcriptomic data for human intestinal enteroids (HIEs) derived from the ileum, jejunum, and duodenum under mock (uninfected) conditions were obtained from publicly available RNA-seq datasets in the NCBI Gene Expression Omnibus (GEO; accession numbers: GSE150918, GSE117911, and GSE183223). Processed transcript abundance values (TPM) provided by GEO were used directly for analysis. No additional normalization or reprocessing was performed.

### Measurement of Caspase-1 Activity

Caspase-1 activity in HIEs following human norovirus infection was measured using the Caspase-Glo® 1 Inflammasome Assay (Promega). HIEs were infected with HNoV in white 96-well plates, and at 48 hours post-infection, Caspase-Glo® 1 reagent was added directly to each well at a 1:1 ratio with the culture medium, according to the manufacturer’s instructions. Plates were incubated for 2 hours at room temperature before luminescence measurement. Luminescence was then measured using a GloMax plate reader (Promega). Caspase-1 activity was calculated based on the luminescence values, which directly reflect enzymatic activity in the samples

### ELISA for IL-1β and IL-18

IL-1β and IL-18 concentrations in cell culture supernatants were measured using commercial ELISA kits according to the manufacturers’ protocols: IL-18 (#7620; R&D Systems) and IL-1β (#DLB50; R&D Systems). Absorbance was measured at 450 nm using a microplate reader, and cytokine concentrations were calculated from standard curves generated with recombinant standards provided in each kit. All samples and standards were assayed in duplicate.

### Generation of CRISPR/Cas9-mediated NLRP6 gene knockout HIEs

NLRP6 knockout HIEs were generated using a CRISPR/Cas9 lentiviral system. Two guide RNAs targeting NLRP6 (gRNA#1: 5’-GCGCGCCTACCGCTTCGTGA-3’, gRNA#2: 5’-TGCCCGCCGCCCAGTCGTAC-3’) were cloned into the pLV[2CRISPR]-hCas9 vector (VectorBuilder). Lentivirus was produced by co-transfecting this plasmid with pMD2.G and psPAX2 into HEK293T cells, concentrated using Lenti-X concentrator (Takara), and stored in basal medium at -70°C. Lentiviral transduction, selection, and expansion of single-cell dissociated HIEs were performed as previously described(30).

### Statistical analysis

All statistical analyses were performed using GraphPad Prism 10. Data are presented as mean ± SEM unless otherwise indicated. Statistical significance was assessed using an unpaired two-tailed Student’s t-test or one-way ANOVA, as appropriate. P-value less than 0.05 were considered significant. All graphical representations and calculations were generated within the GraphPad Prism 10 software environment.

## Acknowledgements

We would like to express our sincere gratitude to Professor Ji-Man Kang (Department of Pediatrics, Yonsei University College of Medicine) for generously providing the human norovirus stool samples used in this study. We also thank Professor Je Wook Yu (Department of Microbiology and Immunology, Yonsei University College of Medicine) for supplying the HEK293T^ASC+CASP1^ cells used in the in vitro reconstituted NLRP3 inflammasome model, and for his valuable guidance on inflammasome-related experimental techniques. In addition, we thank Professor Yoon Jae Song and Na-Eun Kim, Ph.D. (Department of Life Science, Gachon University) for their invaluable assistance in cloning the human norovirus NS7 gene and generating its mammalian expression vector.

## Supporting information

S1 Data. Raw numerical data supporting figure quantification.

S1 Fig. Original and unprocessed images of the figures.

## Notes

### Competing Interest Statement

The authors have declared no competing interest.

